# OCA-B/Pou2af1 is sufficient to promote CD4^+^ T cell memory and prospectively identifies memory precursors

**DOI:** 10.1101/2022.09.21.508912

**Authors:** Wenxiang Sun, Erik P. Hughes, Heejoo Kim, Jelena Perovanovic, Krystal R. Charley, Bryant Perkins, Junhong Du, Andrea Ibarra, Amber R. Syage, J. Scott Hale, Matthew A. Williams, Dean Tantin

## Abstract

The molecular mechanisms leading to the establishment of immunological memory are inadequately understood, limiting the development of effective vaccines and durable anti-tumor immune therapies. Here we show that ectopic OCA-B expression is sufficient to improve antiviral memory recall responses, while having minimal effects on primary effector responses. At peak viral response short lived effector T cell populations are expanded but show increased *Gadd45b* and *Socs2* expression, while memory precursor effector cells show increased expression of *Bcl2*, *Il7r* and *Tcf7* on a per-cell basis. Using an OCA-B mCherry reporter mouse line, we observe high OCA-B expression in CD4^+^ T_CM_ cells. We show that early in viral infection, endogenously elevated OCA-B expression prospectively identifies memory precursor cells with increased survival capability and memory recall potential. Cumulatively, the results demonstrate that OCA-B is both necessary and sufficient to promote CD4 T cell memory in vivo and can be used to prospectively identify memory precursor cells.

**Significance:** CD4^+^ T cell memory is incompletely defined and memory progenitors difficult to identify. Here, we show that expression of the OCA-B transcription coactivator in CD4^+^ T cells is necessary and sufficient to drive productive memory. Using a novel mCherry reporter mouse line, we show that OCA-B expression enriches for responding effector cells with elevated memory potential. The results show that OCA-B expression in T cells is sufficient to promote CD4^+^ memory formation and marks memory precursor cells.

## Introduction

The ability to respond robustly to secondary antigen exposures (immune memory) is a defining feature of the adaptive immune system and forms the physiologic basis for vaccination (1). Immune memory can also drive autoimmune and antitumor responses (2–5). Strategies that promote memory have the potential to improve both vaccine efficacy and tumor immunotherapy. Central memory T cells (T_CM_), which home to secondary lymphoid organs, are particularly long lived and demonstrate high proliferative capacity upon antigen reencounter.

Multiple transcription factors and epigenetic regulators are known to modulate T_CM_ formation or function, including Id3 (6), Blimp1 (7–9), Suv39h1 (10), T-bet (11, 12), Foxo1 (13, 14), Bcl6 (15, 16), Eomes (17) and TCF1 (18). Many of these factors regulate memory more clearly in CD8^+^ compared to CD4^+^ T cells. In CD4^+^ T cells, the lymphocyte-restricted transcriptional coregulator OCA-B (also known as OBF-1 and Pou2af1) is induced upon T cell antigen stimulation (19, 20). T cells from germline OCA-B deficient mice mount normal primary T cell responses to lymphocytic choriomeningitis virus (LCMV) but show attenuated CD4^+^ memory T cell formation and recall response (21). Elevated *Ocab* (*Pou2af1*) mRNA levels are seen in CD4^+^CD25^lo^ early effector cells that are enriched for memory potential (22). OCA-B is also associated with autoimmunity (23–26).

Here, we show that OCA-B deletion in T cells selectively reduces CD4^+^ T cell memory potential. We show that ectopic OCA-B expression minimally affects peak CD4^+^ T cell responses to LCMV, but is sufficient to expand antigen recall responses at the expense of empty vector (EV)-transduced controls. mRNA expression profiling of effector cells ectopically expressing OCA-B reveals gene expression changes associated with effector and memory properties, signaling and homing. These include global increases in *Socs2* and *Gadd45b*, expansion of short-lived terminal effector T cell populations, and increases in *Tcf7* and *Il7r* in specific memory precursor populations. Using an OCA-B reporter mouse, we show that elevated levels of endogenous OCA-B in effector T cells during the primary response prospectively identifies CD4^+^ T effector cells with enhanced survival and expansion capability.

## Results and Discussion

### OCA-B in T cells is required for robust CD4^+^ T cells memory recall response

We used a conditional *Ocab* (*Pou2af1*) mouse allele crossed to a CD4-Cre driver, which efficiently deletes OCA-B in T cells (26), to study effects of T cell-specific OCA-B loss. We infected *Ocab^fl/fl^*;CD4-Cre and control *Ocab^fl/fl^* mice with LCMV^Arm^ and measured specific T cell responses using I-A^b^/gp_66–77_ tetramer staining. No significant differences in the frequency of CD4^+^ I-A^b^/gp_66–77_^+^ CD4^+^ T cells were observed at 8 days post-primary infection (dpi, Fig. S1*A*-*B*, D8). Rechallenging these mice with *Listeria monocytogenes* expressing the LCMV glycoprotein epitope (Lm-gp61) (27) elicited a trending but nonsignificant decrease in the frequency of cells undergoing memory recall response in *Ocab^fl/fl^*;CD4-Cre mice (Fig. S1*A*-*B*, D40+7). The responding cells did however show a significant decrease in IFNγ production (Fig. S1*C*-*D*). The same cells showed minimal changes in ICOS, PD-1, Ly6C or CD62L (Fig. S1*E*). These results indicate that infection of T cell-conditional OCA-B knockout mice with LCMV results in deficiencies in IFNγ expression during memory recall responses.

The relatively small reductions in frequencies of tetramer^+^ CD4^+^ T cells at recall response may reflect selection for a small percentage of cells escaping OCA-B deletion. This selection can be circumvented using bone marrow chimeras in which conditional deficient or control cells are competed against wild-type T cells in the same mice. We generated bone marrow chimeras in which wild-type donors were co-engrafted with congenically-marked *Ocab^fl/fl^*;CD4-Cre or control *Ocab^fl/fl^* cells in the same recipients. After 8 weeks of engraftment, recipient mice were infected with LCMV. The ratio of antigen-reactive *Ocab^fl/fl^* control effector CD4^+^ T cells to wild-type competitor was close to 1:1 at 8 dpi. OCA-B deficient (*Ocab^fl/fl^*;CD4-Cre) cells were slightly reduced relative to wild-type competitor (Fig. S1*F*). To investigate the effects of T cell-specific OCA-B loss on memory recall responses, we rechallenged mice with Lm-gp61. Following rechallenge, the control to wild-type cell ratio remained close to 1:1 at both primary and memory recall response, while wild-type competitor cells were >8-fold increased at the expense of OCA-B deficient cells specifically at recall (Fig. S1*F*-*G*). These findings indicate that, as with T cell-conditional Oct1 deficiency and global OCA-B deficiency (21), T cell-conditional OCA-B deficiency results in severe deficiencies in CD4^+^ T cell memory recall responses.

### Ectopic OCA-B expression is sufficient to enhance CD4^+^ T cell memory responses

To study the effect of forced OCA-B expression, we primed Ly5.1**^+^**/5.2**^+^** (CD45.1**^+^**45.2**^+^**) SMARTA TCR transgenic donor mice with GP_61-80_ peptide and transduced isolated splenic CD4^+^ T cells ex vivo with a retroviral vector expressing mouse OCA-B and GFP. In parallel, Ly5.1**^+^**/5.1**^+^**control SMARTA cells were transduced with GFP-expressing EV. SMARTA mice express a transgenic TCR recognizing an immunodominant LCMV glycoprotein (GP_61–80_) CD4^+^ epitope (28). GFP^+^ SMARTA cells expressing OCA-B and EV controls were sorted, mixed 1:1 and adoptively co-transferred into Ly5.2^+^/5.2^+^ recipients (Fig. 1*A*). Efficient OCA-B expression was confirmed in primary CD4^+^ T cells (Fig. 1*B*). After transfer, recipient mice were infected with LCMV and the cell populations were detected by flow cytometry using congenic markers from the donor T cells and GFP expressed from the viral vector. At 8 dpi, splenic effector EV and OCA-B transduced cells expanded equivalently (Fig. 1*C* and D). Another group of mice were infected with LCMV and allowed to clear the virus and form memory. Recall responses were induced by re-challenge using Lm-gp61. At static memory timepoints (40 dpi) slight but statistically nonsignificant skewing towards the OCA-B population was noted, however at peak T cell response to rechallenge (40+7 dpi), ∼50-fold more OCA-B overexpressing SMARTA cells were present (Fig. 1*C*-*E* and Fig. S2*A*). Although memory recall potential was improved, increases in Tbet and Granzyme B (Gzmb), and decreases in CXCR5, Bcl6 and TCF1 at 8 dpi were consistent with an increased Th1 phenotype in OCA-B transduced cells (Fig. 1*F*, *G* and Fig. S2*B*). No significant differences in IFNγ and Ki-67 were observed at 8 dpi (Fig. 1*H* and *I*). Additionally, IL-7R MFI was significantly decreased in OCA-B expressing cells although cells with strong expression could still be identified (Fig. S2*C*). The ability of ectopic OCA-B to promote memory recall responses was also observed at longer timepoints in both spleen as well as peripheral blood (Fig. J and K and Fig. S2*D*). These results indicate that OCA-B expression enhances the ability of CD4^+^ T cells to induce memory responses.

**Fig. 1.**
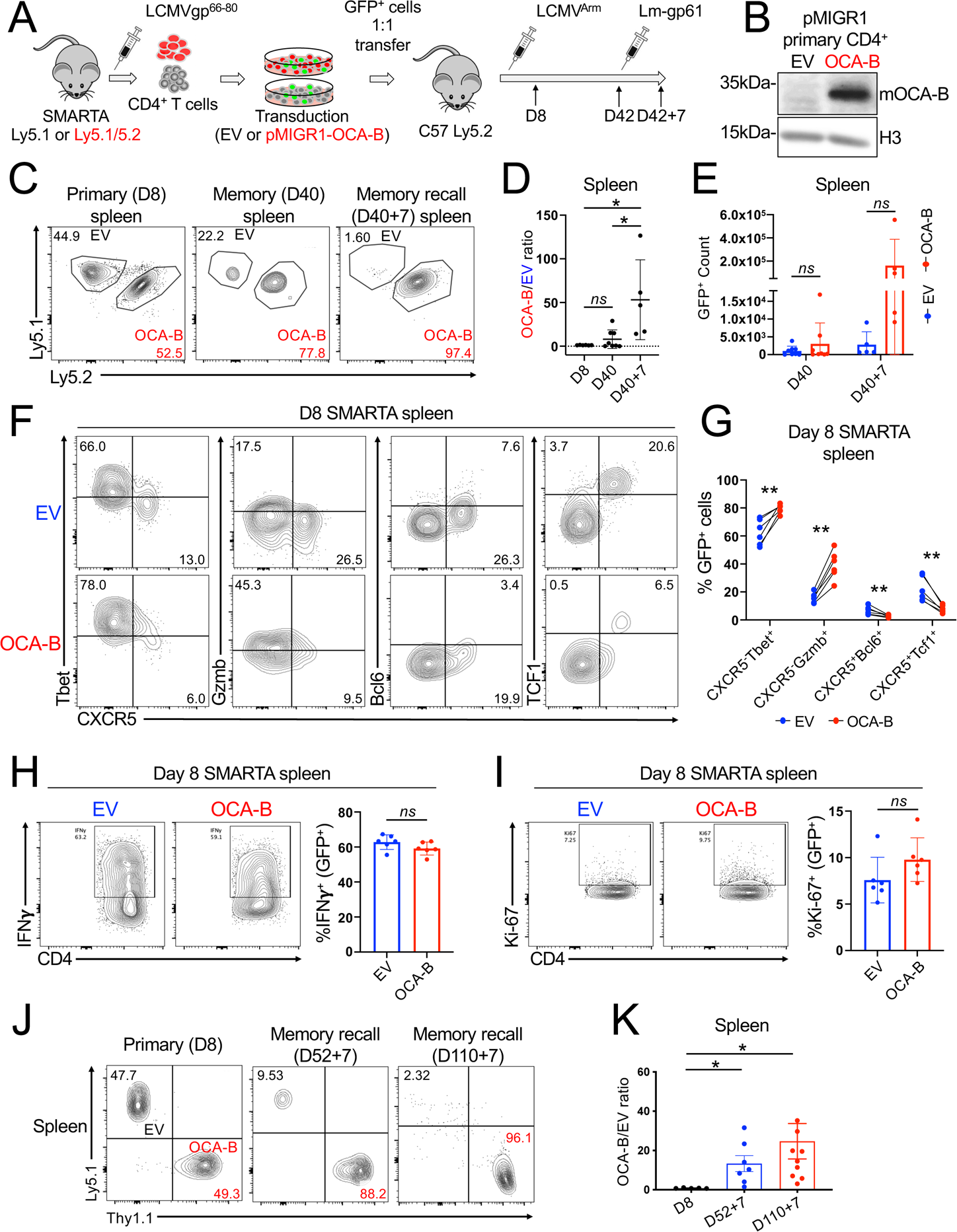
Ectopic OCA-B expression enhances CD4+ T cell memory recall responses in vivo. (*A*) Experimental schematic for OCA-B transduction and T cell mixed adoptive transfer. Ly5.1^+^ SMARTA cells were transduced with pMSCV-IRES-GFP (pMIGR1, EV) and Ly5.1^+^/5.2^+^ SMARTA cells were transduced with pMIGR1 expressing mouse OCA-B (pMIGR1-OCA-B). The donors were age- and sex-matched littermates. Two days following transduction, GFP^+^Ly5.1^+^ SMARTA cells (EV) and GFP^+^Ly5.1^+^/5.2^+^ SMARTA cells (OCA-B) were sorted, combined 1:1, co-transferred into Ly5.2^+^/5.2^+^ C57BL6/J recipient mice, and infected with LCMV. Recipients were tested for static memory on day 40 post-infection, and also rechallenged with Lm-gp61 on day 40 analyzed after 7 days. (*B*) Lysates from primary CD4+ T cells transduced with either empty pMIGR1 (EV) or pMIGR1-OCA-B were immunoblotted using antibodies against OCA-B. Histone H3 is shown as an internal loading standard. (*C*) Flow cytometric analysis of Ly5.1^+^ pMIGR1 EV-transduced and Ly5.1^+^/5.2^+^ pMIGR1-OCA-B-transduced (OCA-B) SMARTA cells in the spleen and blood of a representative recipient mouse at peak effector response (D8), resting memory (D40) and memory recall (D40+7). Live cells were gated based on CD4 and GFP positivity. (*D*) Ratio of OCA-B/EV transduced cells at D8, D40, and D40+7. (*E*) GFP+ cell counts per spleen for EV and OCAB transduced cells D40 and D40+7 after LCMV infection. (*F*) D8 response representative flow cytometry plots showing Tbet, Gzmb, Bcl6, Tcf1, and CXCR5 expression in splenic GFP+ CD4+ T cells by EV or OCA-B transduction condition. (*G*) Mean relative percentages of CXCR5^-^Tbet^+^, CXCR5^-^Gzmb^+^, CXCR5^+^Bcl6^+^, and CXCR5^+^Tcf1^+^ cells between transduction condition at peak response. Individual mice are connected by lines. (*H*) Representative flow plots and frequency quantification of IFNγ producing GFP^+^ EV or OCA-B transduced cells. (*I*) Representative flow plots and frequency quantification of Ki67 producing GFP^+^ EV or OCA-B transduced cells. (J) Representative plots showing relative percentages of splenic EV- or pMIGR1-OCA-B-transduced (GFP^+^) SMARTA cells were plotted at D8, D52+7 and 110+7. Each datapoint represents an individual mouse harboring both EV- and pMIGR1-OCA-B-transduced cells. 5, 7 and 10 mice were sacrificed at each timepoint to obtain measurements from spleen. (*K*) Mean ratios of pMIGR1-OCA-B-transduced relative to EV-transduced SMARTA cells (OCA-B/EV) are plotted. Splenic SMARTA T cells are shown at day 8 post-infection, or 7 days post-rechallenge with Lm-gp61 (52+7 or 110+7).

To investigate if ectopic OCA-B expression induces gene expression changes in effector cells associated with improved memory recall, we profiled gene expression using splenic SMARTA T cells at 8 dpi. CD4^+^ cells were first isolated by magnetic isolation, then congenically marked GFP^+^ cells were further isolated by fluorescence-activated cell sorting (FACS). We identified ∼600 down-regulated genes and ∼200 up-regulated genes (Table S1). Unsupervised hierarchical clustering identified differentially expressed gene clusters (Fig. 2*A*). Down-regulated genes included *Il7r* and *Bcl6*, consistent with the flow cytometry findings at this timepoint. The top up-regulated gene, *Pou2af1*, encodes OCA-B, providing a quality control. Other up-regulated genes were associated with effector activity (*Tbx21*, *Zeb2*), exhaustion (*Ikzf2*/*Helios*) and attenuation of T cell signaling and proliferation (*Socs2*, *Gadd45b*). *Gadd45b* promotes autoimmune disease and anti-tumor immune responses (29–32). Example genome tracks are shown in Fig. 2*B*. Gene ontology (GO) analysis of the up-regulated genes identified terms associated with H3K9me3 regulation. Down-regulated genes by contrast were strongly associated with H3K27me3 in CD4^+^ and CD8^+^ memory T cells (Fig. 2*C*). ChIP-X Enrichment Analysis (ChEA) (33) identifies Tp53 and Smc3 as factors that also regulate the gene set induced by OCA-B ectopic expression (Fig. 2*D*). p53 and SMC3 may therefore also control expression of these genes or could themselves be controlled by OCA-B to regulate gene expression. Sets of significant Molecular Function, Epigenomic Roadmap and ChEA GO terms are shown in Table S2.

**Fig. 2.**
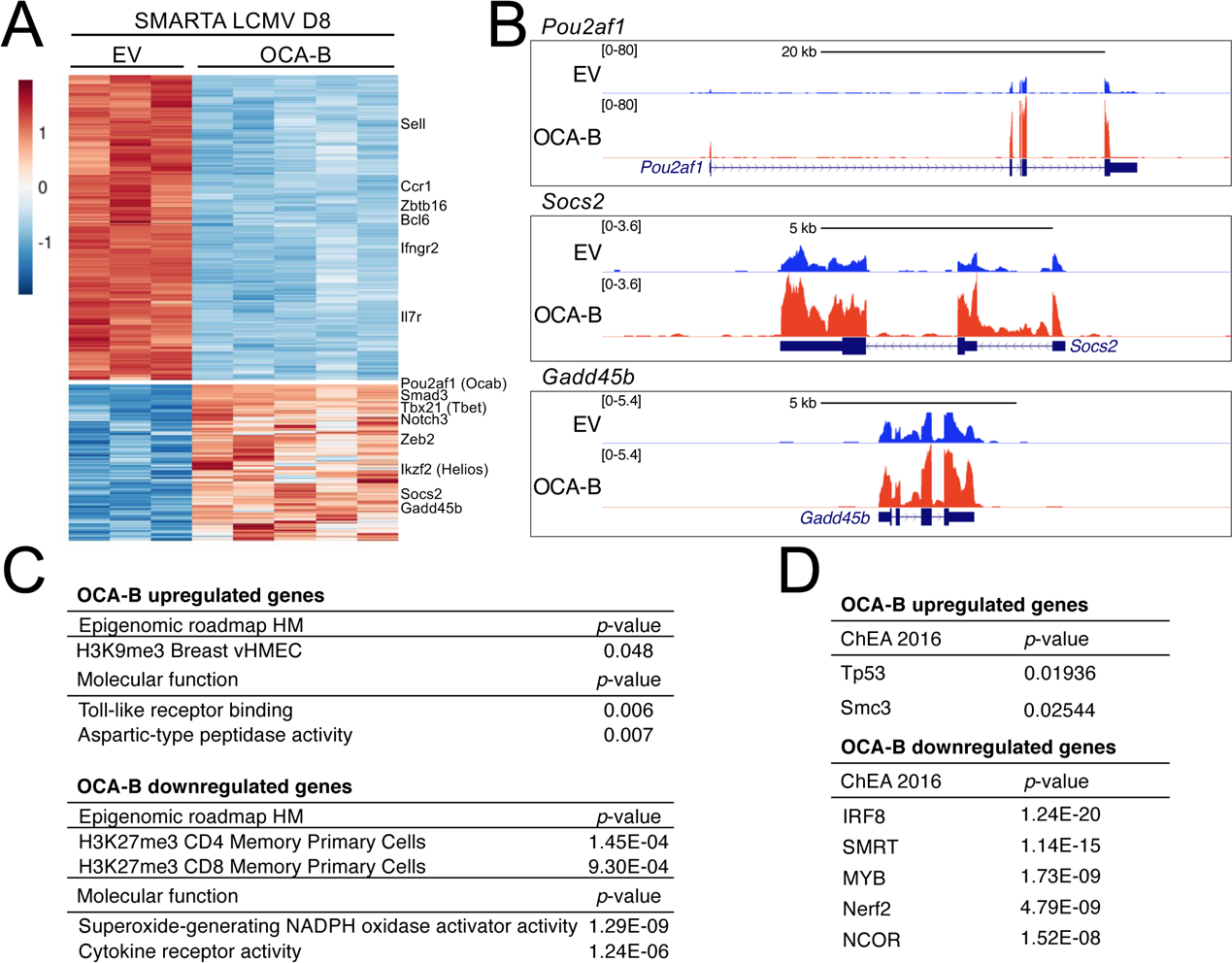
Gene expression changes associated with ectopic OCA-B expression in SMARTA transgenic primary effector T cells. (*A*) Heatmap showing top up- and down-regulated genes. Genes with log2 fold-change >0.5 or <-1.0, and padj ≤ 0.05 are shown. Example genes are shown at right. Control and OCA-B-transduced cells were purified from the same mice using magnetic isolation and FACS. (*B*) Example genome tracks showing two up-regulated genes, *Socs2* and *Gadd45b*. *Pou2af1* (*Ocab*) is shown as a positive control. (*C*) Top Epigenomic roadmap and Molecular function GO terms for the set of OCA-B up- and down-regulated genes. (*D*) Top ChIP-X Enrichment Analysis (ChEA 2016) factors potentially regulating the same set of genes up- and down-regulated by OCA-B ectopic expression.

Effector CD4^+^ T cell populations are heterogeneous, with memory progenitor cells comprising a small proportion of the effector T cell population (12, 22, 34). In bulk RNA-seq, increased expression of genes that promote memory in rare memory progenitors may therefore be obscured by decreases in larger populations of terminal effectors. To test this, we performed single-cell RNA-sequencing (scRNA-seq) at D8 post-LCMV infection. Uniform manifold approximation and projection (UMAP) using EV-transduced cells revealed a variety of populations (Fig. 3*A*, Table S3). Clusters 1 and 4 were enriched for genes such as *Id3*, *Ccr7*, *Bcl2*, *Slamf6* and *Tcf7* (both clusters), *Tox* (cluster 1), and *Cxcr5*, *Il7r*, *Icos* and *Cd69* (cluster 4, Table S3). These clusters likely contain memory precursor effectors. Clusters 0, 2 and 6 by contrast were enriched in *Ccl5*, *Ifng* and *Tbx21*, respectively (Table S3), and likely comprise more terminally differentiated effectors. Cluster 5 was enriched for genes encoding histones, cyclin-dependent kinases and proliferating cell nuclear antigen (PCNA) and likely represent proliferating effectors.

**Fig. 3.**
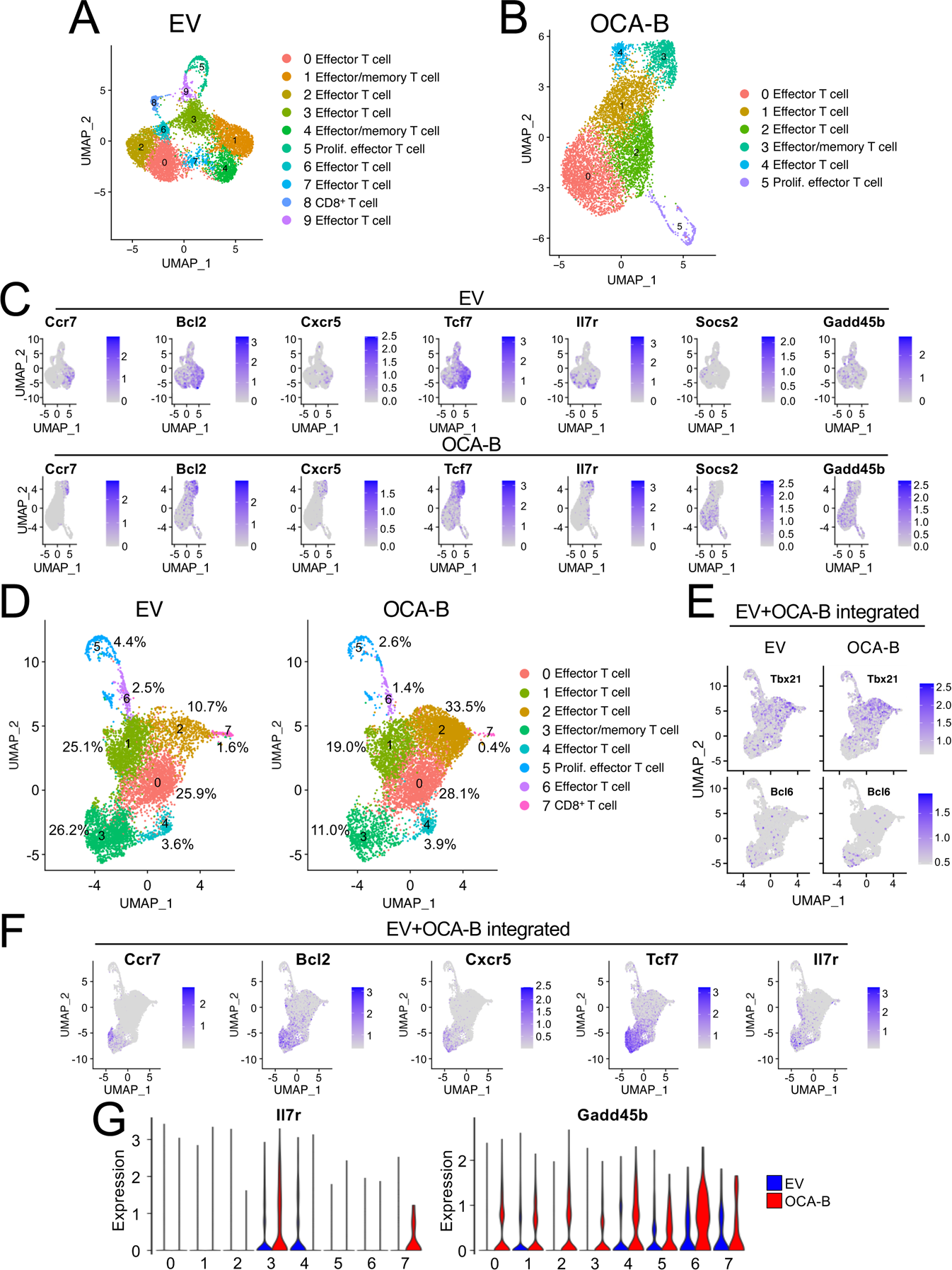
Single-cell RNA-seq analysis of OCA-B-transduced vs. control SMARTA cells at 8 days post LCMV infection. (*A*) UMAP projections of independently clustered EV-transduced (GFP^+^) SMARTA cells purified similar to Figs. 2 and 3, and subjected to scRNA-seq. Cluster identities were annotated using a combination of gene expression enrichment (Table S3), cluster identity predictor ImmGen identity scores (Table S4) and PanglaoDB annotation terms. Clusters 1 and 4 were associated with memory formation. Top PanglaoDB annotation terms for cluster 4 are shown below. (*B*) Similar UMAP projection using OCA-B-transduced cells purified from the same mice as in (A). Cluster identities were annotated similarly to (A). Cluster 3 in OCA-B-transduced cells was most strongly associated with memory formation. Top PanglaoDB annotation terms for cluster 3 are shown below. (*C*) Feature plots highlighting expression of representative genes for the two UMAP projections in (A) and (B). (*D*) UMAP projections of the the EV- and OCA-B-transduced single-cell RNA-seq datasets clustered together. For each projection, the percentage of each cluster relative to the total number cells in condition is shown. Cluster identities were annotated using a combination of gene expression enrichment (Table S3), cluster identity predictor ImmGen identity scores (Table S5) and PanglaoDB annotation terms. (*E*) Feature plots highlighting expression of two representative genes (*Tbx21* and *Bcl6*) associated with specific clusters. (*F*) Additional combined feature plots supporting the association of cluster 3 with memory progenitors. (*G*) Violin plots depicting expression of six genes (*Il7r*, *Zeb2*, *Gadd45b*, *Foxo1*, *Tcf7* and *Tox*) across each of the 10 clusters in EV-transduced cells (blue, left) and OCA-B-transduced cells (red, right).

OCA-B-transduced cells clustered more uniformly with fewer clusters (Fig. 3*B*). Cluster 0 effector cells were marked by *Zeb2*, *Klrg1* and *Tbx21* (Table S3). Clusters 1 and 2 were also associated with effector activity. OCA-B transduced cluster 3 was associated with effector/memory progenitor activity similar to EV-transduced clusters 1 and 4 (Fig. 3*C*). Although there were proportionately fewer of these cells compared to the EV-transduced clusters 1 and 4, multiple memory-associated genes were increased on a per-cell basis. *Tcf7* was increased by >4-fold in OCA-B cluster 3 relative to other clusters, but only 2-fold in EV clusters 1 and 4 (Table S3). *Slamf6* was increased by >3-fold in OCA-B cluster 3, but <2.5-fold in EV clusters 1 and 4. *Bcl2* and *Il7r* were similarly more strongly expressed relative to the other clusters in OCA-B-compared to EV-transduced cells. Additionally, *Foxo1* was not over-represented in either memory-associated EV cluster but was enriched in OCA-B cluster 3, while *Ikzf2*, which has been associated with CD4^+^ T cell exhaustion (35), was not enriched in cluster 3 but was enriched in the equivalent EV-transduced clusters (Table S3). *Id3* was enriched in OCA-B-transduced cluster 3 more than EV cluster 4 but less so than in EV cluster 1. These findings suggest that OCA-B transduced cells in cluster 3 are qualitatively superior to their EV-transduced counterparts at forming memory. Simultaneously, OCA-B-transduced clusters showed broad increases in the expression of *Socs2*, which encodes an attenuator of T cell signaling, and *Gadd45b*, which promotes growth arrest (Fig. 3*C*). These changes likely attenuate effector function and/or may enable a broader transition of effectors to memory progenitors.

Clustering EV- and OCA-B-transduced cells together allows the relative proportions of cells with common gene expression features to be determined. Again, specific clusters were enriched for terminal effectors (Fig. 3*D*, clusters 0, 1, 2, 4, 6), proliferating effectors (cluster 5) and effector cells with memory progenitor potential (cluster 3). Feature plots for genes associated with effector cells (*Tbx21*) and memory function (*Bcl6*) are shown in Fig. 3*E*. Additional plots for other genes associated with memory are shown in Fig. 3*F*. In the OCA-B-transduced condition, cluster 3 contained proportionately fewer cells but showed elevated expression of genes such as *Il7r* and *Gadd45b* on a per-cell basis (Fig. 3*G*). These findings identify changes in gene expression in effector T cell populations consistent with increased memory potential.

### OCA-B knock-in reporter mice label CD4^+^ memory T cell populations

To monitor OCA-B expression in viable cells, we knocked 3 mCherry reporter cassettes preceded by P2A elements into the mouse *Pou2af1* (*Ocab*) locus. The resulting reporter allele contained the mCherry cassettes inserted immediately before the stop codon and the 3’ UTR (Fig. 4*A*). A successful hemizygous knock-in founder and a homozygote generated by intercrossing the progeny were confirmed by PCR (Fig. S3*A*). Hemizygous and homozygous reporter mice retained normal OCA-B protein expression (Fig. S3*B*). OCA-B was readily detectable in peripheral blood B cells from hemizygous and homozygous reporter mice (Fig. S3*C*, *D*). Peripheral T cell expression was weaker, forming a shoulder in hemizygous reporters, but an observable peak in homozygotes (Fig. S3*C*, *D*). We therefore used heterozygous mice to perform baseline characterization of B cell populations, and homozygotes to study T cells.

**Fig. 4.**
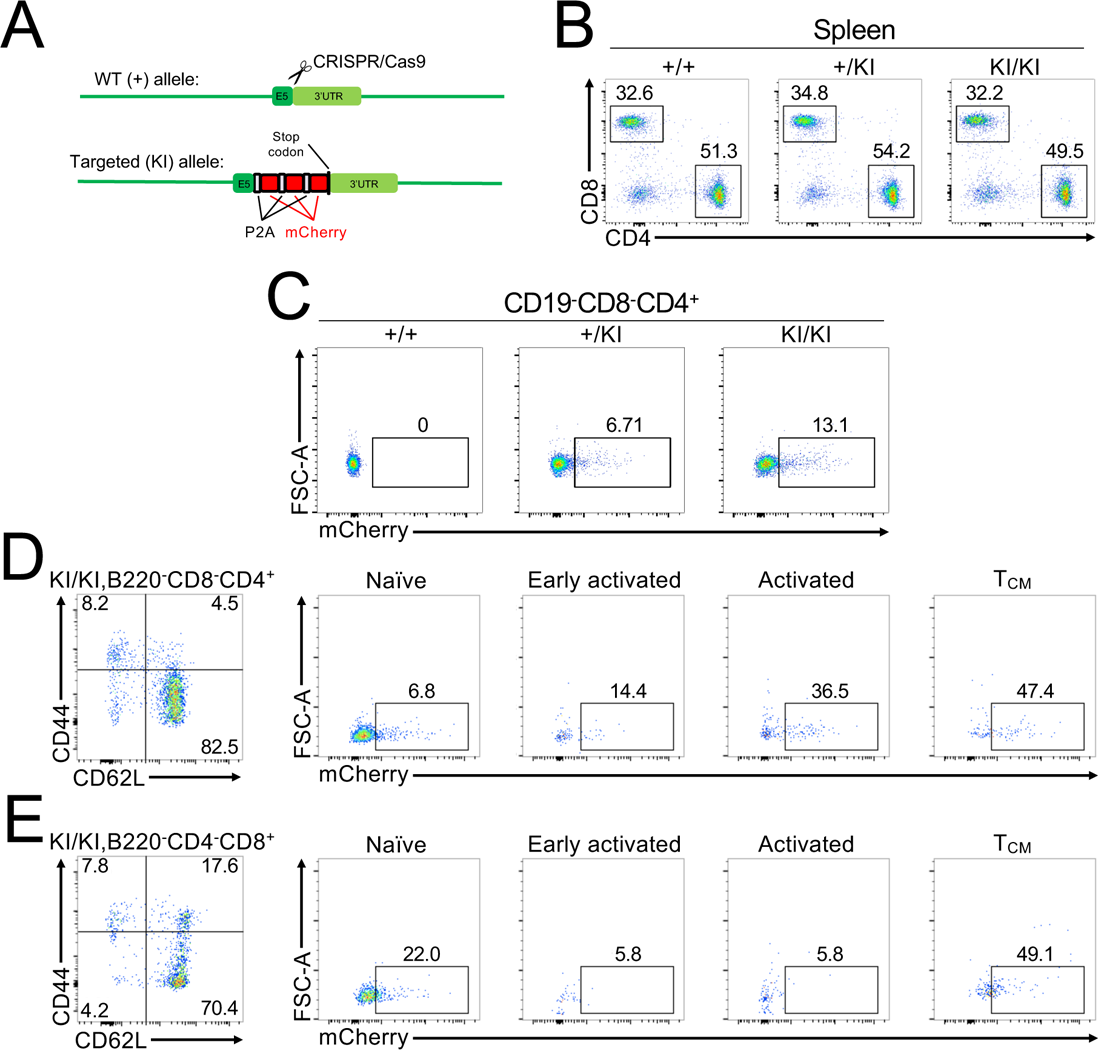
mCherry expression in splenic T cell populations in specific pathogen-free homozygous OCA-B-3×mCherry mice. (*A*) Mouse *Pou2af1* locus, targeting vector and the targeted *Pou2af1* locus. (*B*) CD4 and CD8 expression are shown for an allelic series of example wild-type (+/+) and homozygous (KI/KI) OCA-B knock-in reporter mice. (*C*) OCA-B homozygous reporter expression in splenic CD4^+^ T cell populations. (*D*) The same CD4^+^ cells as in (C) were further stratified by CD62L and CD44 into naïve, early activated, activated and central memory (Tcm) populations. CD62L^lo^CD44_lo_ cells are likely recently activated cells that have down-regulated CD62L but not yet up-regulated CD44. (*E*) Similar analysis as in (D) except for CD8^+^ cells.

Robust mCherry expression could be detected in the bone marrow, peritoneal cavity, spleen and lymph nodes of specific pathogen-free hemizygous reporter mice. Total thymocytes by contrast expressed little OCA-B (Fig. S3*E*). OCA-B expression was readily detected in developing, immature and recirculating B cell populations (Fig. S3*F*). Expression was nearly uniform in the different populations, except for a small population of immature B cells that lacked OCA-B expression. Peritoneal B-1 and B-2 B cells were also homogeneous, with stronger expression in B-1 relative to B-2 cells (Fig. S3*G*). Splenic B cells can be partitioned into transitional and mature populations. T1/2/3 subsets uniformly expressed OCA-B (Fig. S3*H*). Marginal zone (MZ), follicular and non-follicular populations all expressed OCA-B strongly (Fig. S3*I*). Expression was largely uniform, but there was a significant population of newly formed CD21^lo^CD23^lo^ B cells expressing no OCA-B. The nature and significance of these cells is unknown. These results document strong reporter activity in B cell populations, with the highest level of activity in MZ and B-1 B cells. MZ B cells are largely absent in OCA-B knockout mice (36). These results indicate that the OCA-B reporter robustly labels B cells.

Gating out B cells, approximately 5% of thymocytes from unchallenged, specific pathogen-free homozygous reporter mice expressed OCA-B (Fig. S3*J*). mCherry expression could be identified in CD4/CD8 double-negative (DN), DP and SP compartments, with the highest amounts in DN and CD8 SP cells (Fig. S3*K*). In the spleen, the relative abundance of CD4^+^ and CD8^+^ T cells was unaltered by the reporter allele (Fig. 4*B*). Splenic CD4^+^ T cells express mCherry at ∼50-fold lower levels compared to B cells and with a wider range of expression (Fig. 4*C*). Peripheral blood T cell mCherry levels varied over a similar range (Fig. S3*C* and not shown).

We used CD62L and CD44 to stratify resting splenic CD4^+^ and CD8^+^ cells into naïve, activated/effector and central memory compartments. Reporter activity could be detected in approximately 7% of naïve (CD62L^hi^CD44^lo^) cells (Fig. 4*D*). The nature of these cells is unknown. Progressively larger fractions of early activated (CD62L^lo^CD44^lo^), activated (CD62L^lo^CD44^hi^) and T_CM_ (CD62L^hi^CD44^hi^) cells expressed mCherry (Fig. 4*D*). Naïve CD8^+^ T cells showed comparatively higher expression levels, but decreased expression in the activated state and similar expression in T_CM_ cells (Fig. 4*E*). Therefore, within the CD4^+^ T cell compartment, increased OCA-B expression as measured by mCherry fluorescence progressively labels greater fractions of naïve, activated and central memory CD4^+^ cells.

### Elevated OCA-B expression prospectively enriches memory precursor CD4^+^ T cells

To assess OCA-B responses during infection, memory formation and rechallenge, we infected reporter mice with LCMV and used MHC tetramers to identify splenic T cells recognizing immunodominant LCMV epitopes. Prior to infection, >90% of cells were CD44^lo^ naïve phenotype. Up to 30% of cells became CD44^hi^ following infection or rechallenge, and a significant number of these were Tet^+^ as expected (Fig. S4*A*, upper plots). A high fraction of activated CD44^hi^ cells expressed mCherry regardless of tetramer status (Fig. S4*A*, lower plots). mCherry was gated relative to a nonreporter control (Fig. S4*B*). Fewer activated CD8^+^ cells expressed mCherry relative to nonreporter controls (Fig. S4*C*,*D*). After LCMV clearance, memory timepoint CD4^+^ cells maintained high mCherry percentages (Fig. S4*A*, D43, 61.6%), though with ∼2-fold lower expression per cell (Fig. S4*E*). Rechallenge with Lm-gp61 to generate CD4^+^ T cell antigen-specific recall responses reduced the percentage of mCherry^+^ cells compared to resting memory, with high-affinity tetramer^+^ cells declining from >60% to <10% mCherry (Fig. S4*A*, D42+7). In contrast, a higher percentage of CD8^+^ reporter T cells expressed mCherry prior to infection. Higher percentages of tetramer^+^mCherry^+^ cells were also present at 5 dpi, but by 8 dpi (peak T cell response) the mCherry percentage was reduced (Fig. S4*B*). Antigen specific memory CD8^+^ T cells also showed reduced mCherry levels relative to CD4^+^ (Fig. S4*B*). We did not collect data for CD8^+^ rechallenge because Lm-gp61 does not express the CD8-dominant epitope recognized by H2-D^b^ LCMV tetramers.

To determine if OCA-B expression in responding CD4^+^ T cells is associated with increased memory potential, we crossed the reporter allele to a SMARTA TCR transgenic background to fix the TCR specificity and to Ly5.1 to track engrafted cells. We transferred cells into recipients, infected with LCMV, isolated mCherry^hi^ and mCherry^lo^ cells at 8 dpi, transferred large numbers of cells (8×10^5^) into naïve secondary recipient mice, and isolated splenic CD4^+^ T cells to assess contraction (Fig. 5*A*). mCherry gating for the cells is shown in Fig. S4*F*. More cells were recovered from mice engrafted with mCherry^hi^ cells, indicating that these cells contracted less than mCherry^lo^ controls (Fig. 5*B*, *C*). Interestingly the cells that did survive showed similar mCherry levels (Fig. 5*D*). These findings indicate that effector CD4^+^ cells expressing higher OCA-B contract less when transferred into naïve animals.

**Fig. 5.**
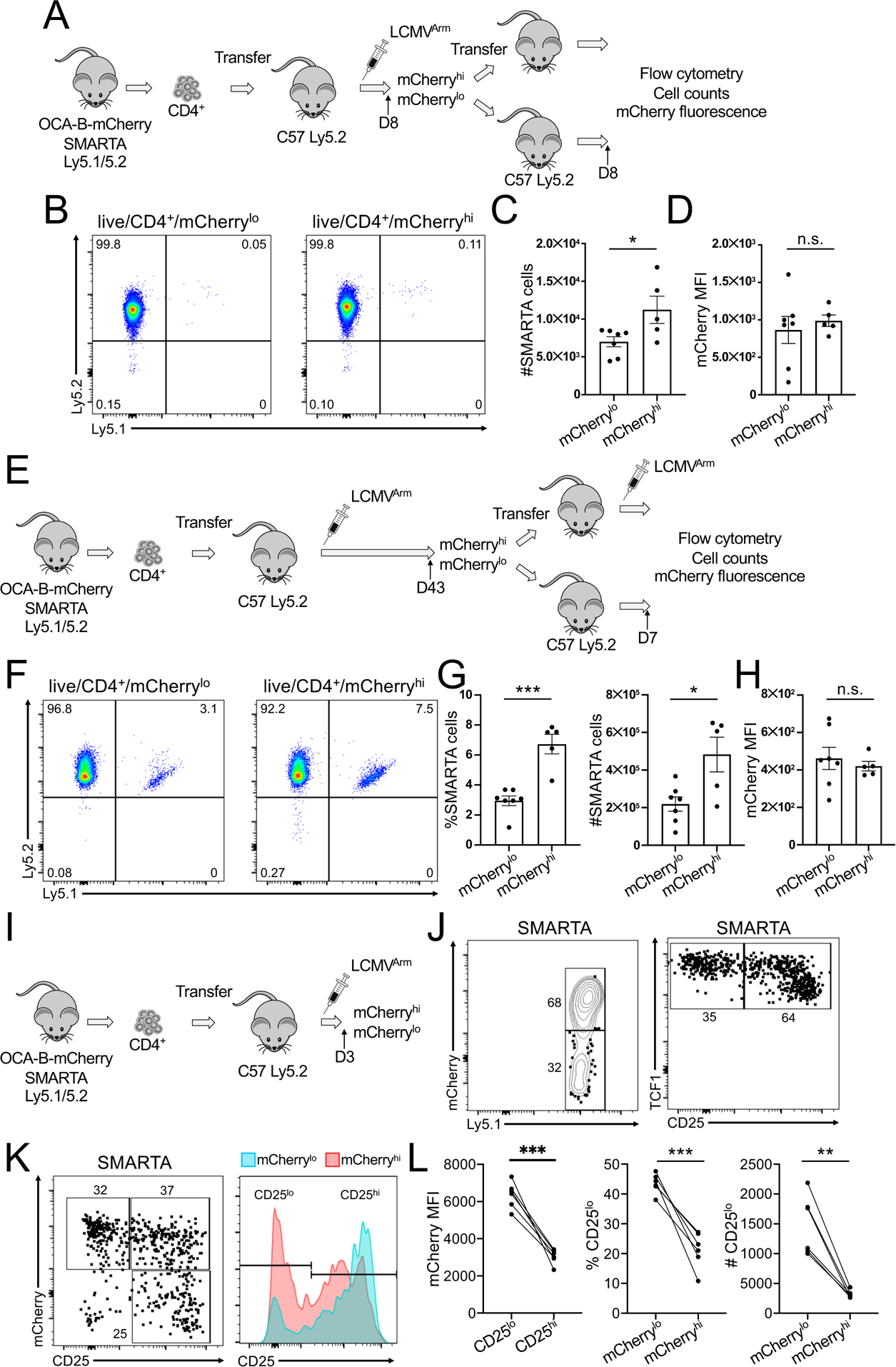
CD4^+^ T cells expressing high levels of OCA-B reporter activity preferentially form central memory cells. (*A*) Schematic for assessing contraction of mCherry^hi^ vs. mCherry^lo^ SMARTA T cells following LCMV infection. Congenically marked SMARTA T cells were isolated from donor mice, transferred into naïve secondary recipients and infected with LCMV. At 8 days post-infection, mCherry^hi^ and mCherry^lo^ populations were isolated by FACS. 8×10^5^ SMARTA cells were transferred into naïve recipients. After 8 days, splenic CD4^+^Ly5.1^+^5.2^+^ SMARTA cells were evaluated by flow cytometry. (*B*) Flow cytometric analysis of SMARTA donor T cells engrafted into naïve mice to monitor rates of decline. Representative mice engrafted with mCherry^lo^ (left) or mCherry^hi^ (right) cells are shown. (*C*) Quantification of averaged donor T cell numbers from mice engrafted with mCherry^lo^ or mCherry^hi^ cells. N=7 for the mCherry^lo^ group and N=5 for the mCherry^hi^ group. (*D*) Similar analysis to (C) except quantifying mean mCherry fluorescence intensity. (*E*) Schematic for assessing recall responses of mCherry^hi^ vs. mCherry^lo^ SMARTA T cells following LCMV infection. Congenically marked SMARTA T cells were isolated from donor mice, transferred into naïve secondary recipients and infected with LCMV. However, mice were allowed to clear LCMV and form memory. After 43 days, mCherry^hi^ and mCherry^lo^ memory T cell populations were isolated by FACS. 1.3×10^4^ SMARTA cells were transferred into naïve recipients, which were infected with LCMV one day later. 7 days post-infection, splenic CD4^+^Ly5.1^+^5.2^+^ SMARTA cells were evaluated by flow cytometry. (*F*) Flow cytometric analysis of SMARTA donor T cells 7 days post-rechallenge to monitor recall responses. Representative mice engrafted with mCherry^lo^ (left) or mCherry^hi^ (right) cells are shown. (*G*) Quantification of averaged donor T cell percentages and numbers from mice engrafted with mCherry^lo^ or mCherry^hi^ cells. N=7 for the mCherry^lo^ group and N=5 for the mCherry^hi^ group. (*H*) Similar analysis to (G) except quantifying mean mCherry fluorescence intensity. (*I*) Schematic for assessing mCherry levels in SMARTA T at early timepoints following LCMV infection. 2×10^5^ congenically marked (CD45.1^+^) SMARTA T cells were isolated from donor mice and transferred into naïve secondary recipients. One day later mice were intraperitoneally infected with 2×10^5^ PFU LCMV. Splenic T cells were collected at 3 dpi. (*J*) Gated unfixed splenic CD4^+^ T cells were assessed for the SMARTA congenic marker Ly5.1 and mCherry (left), while fixed Ly5.1^+^ (SMARTA) cells from the same mice were used to assess CD25 and TCF1 (right). (*K*) CD25 and mCherry expression were assessed from a representative animal. Left: distribution of SMARTA cell CD25 and mCherry expression in SMARTA mice. Right: CD25 levels of gated mCherry^hi^ and mCherry^lo^ SMARTA cells displayed as a concatenated histogram. (*L*) Quantification of averaged mCherry MFI in CD25^hi^ and CD25^lo^ cells (left), and frequencies (center) and numbers (right) of CD25^lo^ cells in mCherry^lo^ or mCherry^hi^ cells SMARTA T cells. N=6 independent recipient mice.

To test the ability of OCA-B expressing cells to mount antigen recall responses, we similarly transferred congenically marked SMARTA cells into recipient mice and infected with LCMV, but collected mCherry^hi^ and mCherry^lo^ cells at 43 dpi. 4×10^4^ memory SMARTA cells were transferred into naïve hosts, which were challenged with LCMV and analyzed at 7 dpi (Fig. 5*E*). mCherry^hi^ SMARTA cells responded better compared to mCherry^lo^ (Fig. 5*F*, *G*) though again with expression normalizing following antigen re-encounter (Fig. 5*H*). Example flow cytometry plots are shown in Fig. S4G.

In mice, OCA-B expression can be detected, varying over a 10-fold range, in responding cells as early as one day after LCMV infection (21). Memory progenitor cells characterized by low CD25 expression can be identified as early as early as 3 dpi (15, 22). We transferred SMARTA homozygous OCA-B mCherry reporter T cells into congenic recipients, infected with LCMV, and collected spleens at 3 dpi to study mCherry and CD25 levels (Fig. 5*I*). A large fraction of responding CD25^lo^ SMARTA cells expressed mCherry (Fig. 5*J*). SMARTA cells from the same mouse expressing low levels of CD25 also expressed high levels of TCF1, as expected (22). mCherry and TCF1 were not compared as cell fixation quenches mCherry, however we were able to directly compare CD25 and OCA-B. CD25^hi^ cells expressed a range of OCA-B, while CD25^lo^ cells were highly enriched for mCherry^hi^ cells (Fig. 5*K*, *L*). These findings show that high OCA-B expression can be used to prospectively mark live CD4^+^ memory precursor effector T cells with augmented capacity to form central memory. Cumulatively, this study indicates that OCA-B expression within CD4^+^ T cells is both necessary for, and sufficient to promote, the emergence of T_CM_.

## Materials and methods

### Mice

All mice used in this study were on the C57BL6/J strain background. All mouse experiments were approved by the University of Utah Institutional Animal Care and Use Committee (protocol 00001553). *Pou2af1* (*Ocab*) conditional (floxed) mice were described previously (26). *Pou2af1*-3×mCherry reporter knock-in mice were generated on a C57BL/6N background (Biocytogen), but were backcrossed >5 times to C57BL/6J prior to generation of data herein.

### LCMV and *Listeria* infection

LCMV Armstrong 53b (LCMV^Arm^) (37) was grown in BHK cells and titered using Vero cells (28). For primary infection, 8-to 12-week-old *Ocab^fl/fl^* or *Ocab^fl/fl^*;CD4-Cre mice were inoculated i.p. with 2×10^5^ plaque-forming units (PFU) of LCMV intraperitoneally.

For heterologous rechallenge with *Listeria monocytogenes* expressing LCMV glycoprotein 61-80 (Lm-gp61) (27), bacteria were grown in log phase in BHI media, and concentrations were calculated using OD at 600 nm (OD of 1=1×10^9^ CFU/mL). Mice were rechallenged intravenously (i.v.) with 2×10^5^ colony forming units (CFU) of Lm-gp61 the indicated number of days after primary infection as published (38).

### Bone marrow radiation chimeras

Radiation chimeras were generated as published (39). Briefly, male Ly5.1 C57BL/6J mice were used as recipients. Mice received a split dose of 900 Rad (2×450 RAD, spaced 1 hr apart), and were engrafted one day later with 1 million mixed donor bone marrow cells. Competitor bone marrow cells from Ly5.2/Thy1.1 but otherwise wild-type mice, and experimental cells from Ly5.2/Thy1.2 *Ocab^fl/fl^* or littermate *Ocab^fl/fl^*;CD4-Cre mice, were isolated from femurs and tibias of donor mice. After red blood cell lysis, CD3^+^ cells were depleted using biotinylated anti-CD3 antibodies (eBioScience) and anti-biotin magnetic MicroBeads (Miltentyi Biotech) according to the manufacturer’s instructions. CD3-depeleted bone marrow cells were mixed 1:1 and injected retro-orbitally. The chimeras were rested for 8 weeks to allow engraftment, after which mice were infected as above. Virus-reactive cells were identified by LCMVgp_66–77_:I-Ab tetramer staining.

### pMSCV-IRES-GFP-OCA-B vector generation

Wild-type *Ocab* cDNA was amplified from the plasmid pcDNA3.1-OCA-B (40) (a gift from Robert Roeder, Rockefeller University) by PCR and cloned into the pMSCV-IRES-GFP (pMIGR1) retroviral vector (a gift from Ryan O’Connell, University of Utah). Primers used for amplification were mOcab-F-XhoI, 5’CTCGAGCTGTCTGCTTCAAAGAGAAAAGGCAAC; mOcab-R-*Eco*RI, 5’ GAATTCCTAAAAGCCCTCCACGGAGAGGGT. The PCR product was gel purified, digested with *Eco*RI and *Xho*I, and inserted into a similarly digested pMIGR1 backbone using T4 DNA ligase to generate pMIGR1-OCA-B. All constructs were validated by resequencing.

### CD4^+^ T cell isolation and stimulation

Naïve splenic CD4^+^ T cells were isolated and simulated as described previously (41). Briefly, spleens were dissociated by grinding and passing through a 70 µm nylon strainer. Red blood cells were lysed by ACK lysis buffer (150 mM NH_4_Cl, 10 mM KHCO_3_, 0.1 mM EDTA). Cells were isolated from mice using a naïve CD4^+^ T cell isolation kit (Miltenyi), and stimulated in culture using 10 μg/mL plate-bound anti-CD3ɛ and 2 μg/mL anti-CD28 antibodies (eBioscience).

### Ectopic OCA-B expression in CD4^+^ T cells

Ly5.1**^+^**/5.1**^+^**SMARTA and Ly5.1**^+^**/5.2**^+^** SMARTA donor mice were i.v. primed with 200 μg GP_61-80_ peptide (AnaSpec). The next day, T cells were purified using a CD4^+^ T cell isolation kit (Miltenyi). Cells were transduced by spin infection with pMIGR retroviruses packaged in 239T cells, with or without mouse OCA-B cDNA. For spin infection, cells were centrifuged with 293T retroviral supernatant at 1000×*g* for 2 hr at 37°C in the presence of 4 µg/mL polybrene (Sigma). Following spin infection, cells were cultured in RPMI 1640 medium supplemented with 10% fetal bovine serum, 50 U/mL penicillin, 50 µg/mL streptomycin, 2 mM L-glutamine, 1 mM sodium pyruvate, 1× MEM nonessential amino acids, 55 µM 2-mercaptoethanol, and 20 IU/mL recombinant human IL-2 for two days. All components were from ThermoFisher with the exception of IL-2, which was supplied by R&D Systems. After two days, transduced GFP^+^ cells were isolated by FACS (BD FACSAria). 1×10^4^ pMIGR1- and pMIGR1-OCA-B-transduced cells were combined and co-transferred into wild-type (Ly5.2/5.2) C57BL6/J mice. Chimeric mice were infected with LCMV and re-challenged with Lm-gp61 as described above. For the experiment shown in Fig. 1*J*, *K*, T cells from Ly5.1 SMARTA and Thy1.1 SMARTA mice were purified using a CD4^+^ T cell isolation kit (Miltenyi) and stimulated with CD3ɛ/CD28 for 2 days followed by spin transfection.

### Flow cytometry

Spleens were dissociated and passed through a 70 µm nylon strainer. Red blood cells were lysed by ACK lysis buffer (150 mM NH_4_Cl, 10 mM KHCO_3_, 0.1 mM EDTA). For intracellular staining, cell suspensions in RPMI supplemented with 10% fetal bovine serum were re-stimulated for 4 hr with 1 μM LCMV GP_61–80_ or 0.1 μM GP_33-41_ peptide along with Brefeldin A (GolgiPlug Becton-Dickinson, 1 μl/mL) as published (42, 43). Cells were subsequently fixed by cell fixation/permeabilization solution (Cytofix/Cytoperm, Becton-Dickinson) according to the manufacturer’s protocol. For tetramer staining, APC-conjugated gp_66–77_:I-A^b^ and gp_33-41_:H-2D^b^ tetramers were provided by the NIH Tetramer Core Facility (Emory Vaccine Center). Cell suspensions were incubated at 37°C for 3 hr in RPMI with the tetramer followed by cell-surface staining. Tetramer fluorescence was normalized with isotype tetramer (hCLIP:I-A^b^) staining. Fig. 1*C, H*, and *I* were generated using a Cytek Aurora spectral cytometer. All other flow cytometry data were generated using a BD Fortessa LSR. The following anti-mouse antibodies were supplied by Ebioscience: CD4-APC (RM4-5), CD3-Biotin (145-2C11), CD90.2-FITC (53-2.1), IL-2-PE (JES6-5H4), IFNy-APC (XMG1.2), CD8a-PerCP/Cy5.5 (53-6.7), CD90.1/Thy1.1-APC (HIS51). The following antibodies were supplied by Biolegend: CD4-PerCP/Cy5.5 (RM4-5), CD44-APC/Cy7 (IM7), CD278/ICOS-BV510 (C398.4A), CD62L-FITC (MEL-14), CD279/PD-1-BV605 (29F.1A12), CD45.1-PerCP-Cy5.5 (A20), CD45.1-BV711 (104), CD45.2-FITC (104), IgM-APC/Cy7 (RMM-1), B220-APC (RA3-6B2), CD21-PE (7E9), CD23-FITC (B3B4), CD93-APC (AA4.1), CD19-FITC (1D3/CD19), Tbet-APC (4B10), CXCR5-PE-Cy7 (L138D7), CD127/IL7Ra-PE/Cy5 (A7R34), IFNy-APC (XMG1.2), CD8a-APC (53-6.7). The following antibodies were supplied by BD Biosciences: Ly-6C-BV450 (AL-21), Bcl6-BV421 (k112-91), Tcf1-PE (S33-966), Ki67-V450 (B56), CD4-BUV395 (GK1.5), CD19-BUV661 (ID3), CD8a-BUV737 (53-6.7). Gzmb-PE (NGZB) was supplied by Invitrogen. Tcf1-AF488 (C63D9) was supplied by Cell Signaling.

### Bulk RNAseq

CD4^+^ T cells were purified from female Ly5.1 SMARTA and Ly5.1/5.2 SMARTA mice, and separately transduced with pMIGR1-OCA-B or pMIGR1 (EV), respectively. On the following day, transduced GFP^+^ cells were sorted, 20,000 of each combined 1:1, and injected intravenously into 24 male C57BL/6J male recipients. 24h later, each recipient mouse was injected intraperitoneally with 2×10^5^ pfu LCMV. After eight days, the 24 mice were divided into six groups of four mice by combining four spleens together. CD4^+^ T cells were purified using a Miltenyi CD4 T cell isolation kit. Cells then were stained with PerCP-Cy5.5 Ly5.1 and AF700 Ly5.2 antibodies. GFP^+^Ly5.1^+^Ly5.2^+^ (EV) and GFP^+^Ly5.1^+^ (OCA-B-expressing) cells were sorted by FACS and used for RNA purification (RNeasy Mini Kit, QIAGEN). RNA concentrations were determined using a Quant-iT RNA assay kit and a Qubit fluorometer (ThermoFisher). Because limited RNA from EV-transduced cells was obtained, the six EV samples were combined into three samples. Five OCA-B-transduced samples (each comprising four mice) were submitted for RNAseq analysis. These showed a high degree of concordance (not shown). Total RNA samples (200-500 ng) were hybridized with Ribo-Zero Plus (Illumina) to deplete cytoplasmic and mitochondrial and ribosomal RNA from samples. RNA sequencing libraries were prepared as described using the Stranded Total RNA Prep, Ligation with Ribo-Zero Plus kit (Illumina) and IDT for Illumina RNA UD Indexes Set A, Ligation (Illumina). Purified libraries were qualified on an Agilent Technologies 2200 TapeStation using a D1000 ScreenTape assay. The molarity of adapter-modified molecules was defined by qPCR using the Kapa Biosystems Kapa Library Quant Kit. Individual libraries were normalized to 0.65 nM in preparation for Illumina sequence analysis. Sequencing libraries were chemically denatured and applied to an Illumina NovaSeq flow cell using the NovaSeq XP workflow (20043131). Following transfer of the flow cell to an Illumina NovaSeq 6000 instrument, a 150 x 150 cycle paired-end sequence run was performed using a NovaSeq 6000 S4 reagent Kit v1.5 (Illlumina).

### Bulk RNA-seq analysis

Bulk RNA-seq analysis was performed as previously described (44). Briefly, Reads were aligned to *Mm10* using STAR (v2.7.3a) and checked for quality using multiqc (v1.10). Between 12 and 14 million pair-end reads were generated for each sample, with >98% of aligned reads mapping to the correct strand and >93% of the reads uniquely aligned to the gene. Differentially expressed genes were identified using DESeq2 version 1.24.0 (45) with a 5% FDR cutoff. Features with zero counts and 5 or fewer reads in every sample were removed from the analysis. Genes increased by 1.4-fold or more or decreased by 2-fold or more, and with adjusted *p*<0.05 were selected as differentially expressed (0.5 < log2FC < -1; padj ≤ 0.05). Figures were generated in R version 4.0.0 using functions from ggplots libraries and pheatmap.

### Single-cell RNA-seq

To profile gene expression at the single-cell level, splenic GFP^+^Ly5.1^+^Ly5.2^+^ (EV-transduced) and GFP^+^Ly5.1^+^ (OCA-B-transduced) SMARTA cells from four pooled mice per condition were isolated, resuspended in PBS with 0.04% bovine serum albumin (ThermoFisher), and filtered through 40 µm strainers. Viability and cell count were assessed using a Countess II (ThermoFisher). Equilibrium to targeted cell recovery of 6,000 cells along with Gel Beads and reverse transcription reagents were loaded to Chromium Single Cell A to form Gel-bead-in Emulsions (GEMs). Within individual GEMs, cDNA generated from captured and barcoded mRNA was synthesized by reverse transcription at 53°C for 45 min. Samples were then heated to 85°C for 5 min. Single cell transcriptomes were assessed using a 10X Genomics Chromium Single Cell Gene Expression instrument. Individual cells were tagged with 16 bp barcodes, and specific transcripts with 10 bp Unique Molecular Identifiers (UMIs) according to manufacturer instructions.

### Single-cell RNA-seq analysis

Single cell transcriptome data were analyzed and clustered as described previously (44). Sequences from the Chromium platform were de-multiplexed and aligned using CellRanger ver. 3.1.0 (10X Genomics) with default parameters mm10-3.0.0. Clustering, filtering, variable gene selection and dimensionality reduction were performed using Seurat ver.4.0.4 (46) according to the following workflow: 1, Cells with <200 detected genes were excluded further analysis. 2, Cells with <5% UMIs mapping to mitochondrial genes and *Cd44* expression >0 were retained for downstream analysis, as SMARTA cells are uniformly responding to LCMV glycoprotein antigen at this time. 3, The UMI counts per ten thousand were log-normalized for each cell using the natural logarithm. 4, Variable genes (2000 features) were selected using the FindVariableFeatures function. 5, Common anchors between the conditions were identified using FindIntegrationAnchors function that were further used to integrate these sets. 6, Gene expression levels in the integrated set were scaled along each gene and linear dimensional reduction was performed. The number of principal components was decided through the assessment of statistical plots (JackStrawPlot and ElbowPlot). 7, Cells were clustered using a shared nearest neighbor (SNN) modularity optimization-based clustering algorithm and visualized using two-dimensional uniform manifold approximation and projection (UMAP). 8, One cluster in each condition marked predominantly by mitochondrial genes (indicative of dying cells) was excluded from the analysis. This resulted in 12,886 EV-transduced cells with 45,748 mean reads per cell, and 2,595 median genes per cell. The total number of reads was 589,513,069, with 59.6% mapping to exonic regions. Similarly, there were 11,709 OCA-B-transduced cells with 68,931 mean reads per cell, and 2,668 median genes per cell. Total number of reads was 807,111,486, with 57.4% mapping to exonic regions. Clusters were identified using manual interrogation of gene expression enrichment for each population (Table S3), the ImmGene cluster identity predictor (Table S4, S5), and PanglaoDB annotation terms. Clusters comprised of contaminating macrophages and neutrophils were manually deleted from the visualization.

### Memory progenitor isolation, normalization and transfer

To assess contraction of the effector cells with different OCA-B levels, splenic CD4^+^ T cells were purified from 8 week-old female Ly5.1/5.2 homozygous knock-in reporter, SMARTA TCR transgenic mice. 2×10^4^ SMARTA cells were intravenously injected into 15 male Ly5.2/5.2 C57BL/6J recipients, which were infected intraperitoneally 1 day later with 2×10^5^ plaque-forming units of LCMV in 250 µL phosphate-buffered saline. After 8 days, responding Ly5.1^+^5.2^+^ SMARTA T cells were sorted into mCherry^hi^ and mCherry^lo^ populations. Cells were pooled and 8×10^5^ mCherry^hi^ and mCherry^lo^ cells were separately injected intravenously into 7 naïve age-matched 9 week-old male C57BL/6J secondary recipients in the case of mCherry^lo^ and 5 secondary recipients in the case of mCherry^hi^. After 8 additional days, mice were sacrificed and splenic Ly5.1^+^5.2^+^ SMARTA cells were counted by flow cytometry. To assess proliferative responses to rechallenge, 15 similar virus-naïve secondary recipients were rested for 43 days post-infection prior to isolation of mCherry^hi^ and mCherry^lo^ populations. 1.3×10^4^ SMARTA cells were transferred into naïve recipients, which were infected with LCMV one day later. 7 days post-infection, splenic CD4^+^Ly5.1^+^5.2^+^ SMARTA cells were evaluated by flow cytometry. Mice with poor infection/response (total splenocyte count <6×10^7^) were excluded from the analysis.

### Quantification and statistical analysis

Excel (Microsoft) and Prism (GraphPad) were used for statistics and graphing. Two-tailed Student T-tests were used to ascribe statistical significance unless otherwise indicated. For all figures, *=*p*-value≤0.05; **=*p*≤0.005; ***=p<0.001. All error bars denote ± standard error of the mean.

## Acknowledgements

We thank M. Chandrasekharan for help with graphics. We thank J. Marvin and the University of Utah Health Sciences Center Flow Cytometry Core facility. We thank O. Allen, B. Dalley and the High-Throughput Genomics Core. MHC tetramers were provided by the NIH Tetramer Core Facility (Atlanta, GA). We thank R. Roeder for the mouse OCA-B cDNA and R. M. O’Connell for the pMIGR1 plasmid.

## Data and materials availability

All data are available in the main text or the supplementary materials. Materials used in the study are available to any researcher for purposes of reproducing or extending the findings. Bulk and single-cell RNA-seq data have been deposited at the Gene Expression Omnibus GEO) website (https://www.ncbi.nlm.nih.gov/geo/, GSE214310).

## Author contributions

J.S.H, M.A.W., D.T. designed research; W.S., E.P.H., H.K., K.R.C., B.P., J.D., A.I. performed research; J.S.H., M.A.W. contributed research tools; W.S., E.P.H., H.K., J.P., K.R.C., B.P., A.R.S analyzed data; W.S., E.P.H., H.K., J.P., K.R.C., J.D., A.I., J.S.H., M.A.W., D.T. wrote the manuscript.

## Classification

Immunology

