## Supplementary material for "OCA-B/Pou2af1 is sufficient to promote CD4^+^ T cell memory and prospectively identifies memory precursors": Supplemetary Figs. S1-S4

### Transcription coactivator OCA-B/Pou2af1 is sufficient to promote T cell-intrinsic CD4 memory and prospectively identifies memory precursors

Wenxiang Sun, Erik P. Hughes, Heejoo Kim, Jelena Perovanovic, Krystal R, Charley, Bryant Perkins, Junhong Du, Andrea Ibarra, J. Scott Hale, Matthew A. Williams, Dean Tantin

Supplementary Information Appendix

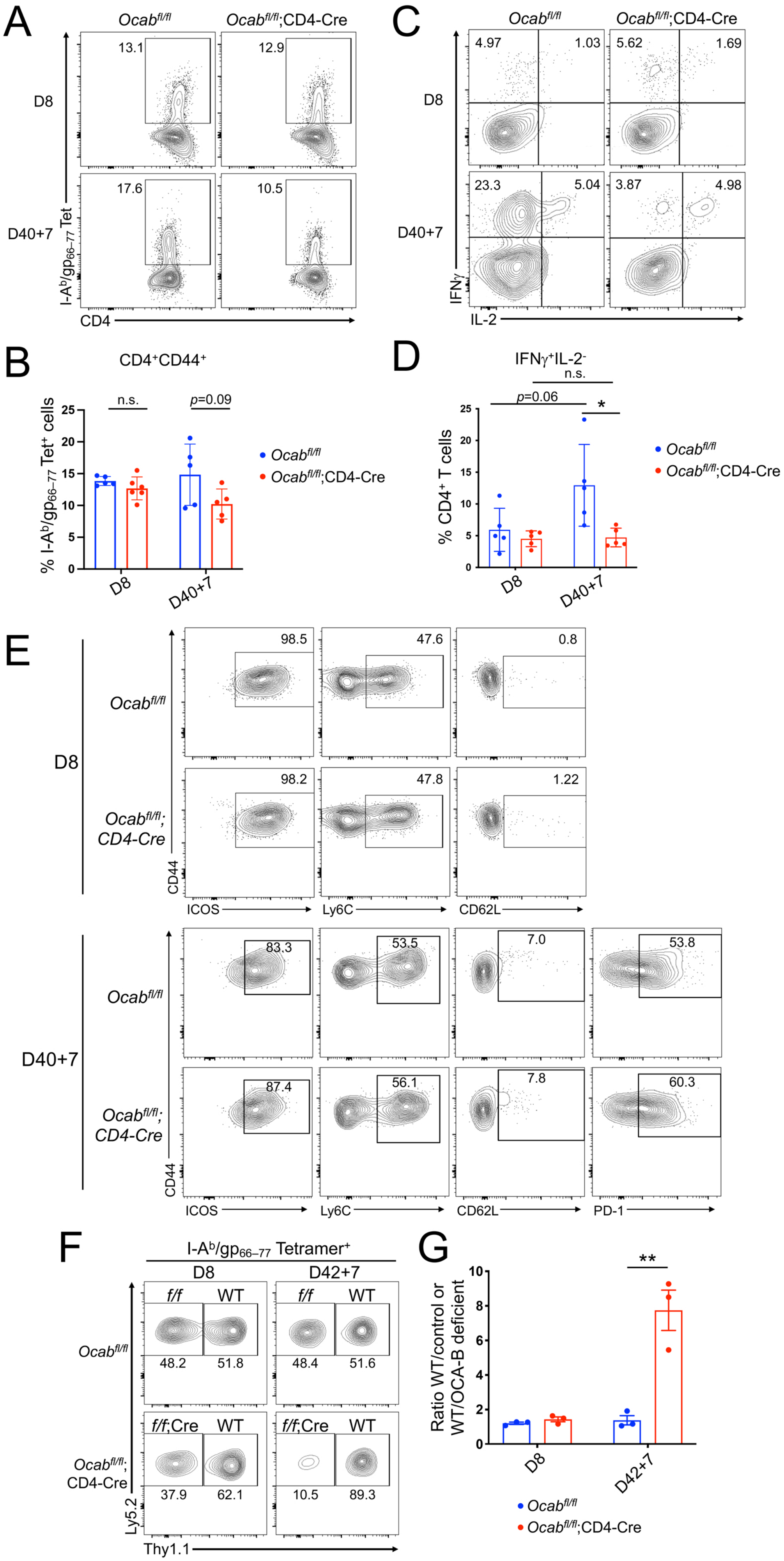
A
Ocabfl/fl
Ocabfl/fl;CD4-Cre
D8
D40+7
13.1
12.9
17.6
10.5
I-Ab/gp66–77 Tet
CD4
C
4.97
1.03
5.62
1.69
23.3
5.04
3.87
4.98
IFNγ
IL-2
B
CD4+CD44+
n.s.
p=0.09
% I-Ab/gp66–77 Tet+ cells
D
IFNγ+IL-2-
n.s.
p=0.06
*
% CD4+ T cells
E
98.5
47.6
0.8
98.2
47.8
1.22
83.3
53.5
7.0
53.8
87.4
56.1
7.8
60.3
CD44
ICOS
Ly6C
CD62L
PD-1
F
I-Ab/gp66–77 Tetramer+
D42+7
f/f
WT
f/f;Cre
48.2
51.8
48.4
51.6
37.9
62.1
10.5
89.3
Ly5.2
Thy1.1
G
**
Ratio WT/control or WT/OCA-B deficient
Ocabfl/fl;CD4-Cre

**Fig. S1.** OCA-B in T cells is required for robust polyclonal memory CD4+ T cell formation and cytokine expression following LCMV infection. (*A*) *Ocab*$^{fl/fl}$;CD4-Cre and control *Ocab*$^{fl/fl}$ mice were infected with LCMV and rechallenged with Lm-gp61 40 days after LCMV infection. Splenic I-A$^b$/gp$_{66–77}$ tetramer-positive, CD4$^+$CD44$^+$ T cells were analyzed by flow cytometry 8 days after primary infection and 7 days after heterologous rechallenge, representative flow cytometry plots are shown. (*B*) Mean T cell frequencies of CD4$^+$CD44$^+$ I-A$^b$/gp$_{66–77}$ tetramer-positive cells are shown. n=5 for each group. n.s.=not significant. (*C*) Splenocytes from LCMV-infected *Ocab*$^{fl/fl}$;CD4-Cre or control *Ocab*$^{fl/fl}$ mice were restimulated in vitro with gp$_{61-80}$ peptide for 4 hours, and IFNγ and IL-2 expression in CD4$^+$ T cells examined by flow cytometry. Cells were gated on CD4 and CD44 positivity. Data from representative mice are shown. (*D*) Mean T cell frequencies of CD4$^+$CD44$^+$IFNγ$^+$IL-2$^-$ T cells are shown. n=5 for each group. (*E*) Surface marker expression in polyclonal OCA-B T cell-deficient and control splenic CD4$^+$ effector T cells at 8 days post-infection or 7 days post-rechallenge. Example flow cytometry panels of ICOS, Ly6C, CD62L. PD-1 was additionally performed for memory recall responses. Cells were gated for I-A$^b$-LCMV$^{gp66-77}$ positivity. (*F*) Flow cytometry plot showing Thy1.1$^+$Ly5.2$^+$ but otherwise genetically wild-type (WT) donor bone marrow cells engrafted 50:50 with Ly5.2$^+$ control cells lacking the Cre driver (*fl/fl*) or experimental T cell-conditional *Ocab*$^{fl/fl}$;CD4-Cre donor cells (*fl/fl*;Cre). Recipient hosts were Ly5.1$^+$. Splenic effector cells were recovered at primary response (D8) or seven days post-rechallenge (D42+7). Cells were gated based on I-A$^b$/gp$_{66–77}$ tetramer positivity. Example mice are shown. (*G*) Mean wild-type competitor to control of OCA-B deficient T cell ratios are shown. n=3 for each group.

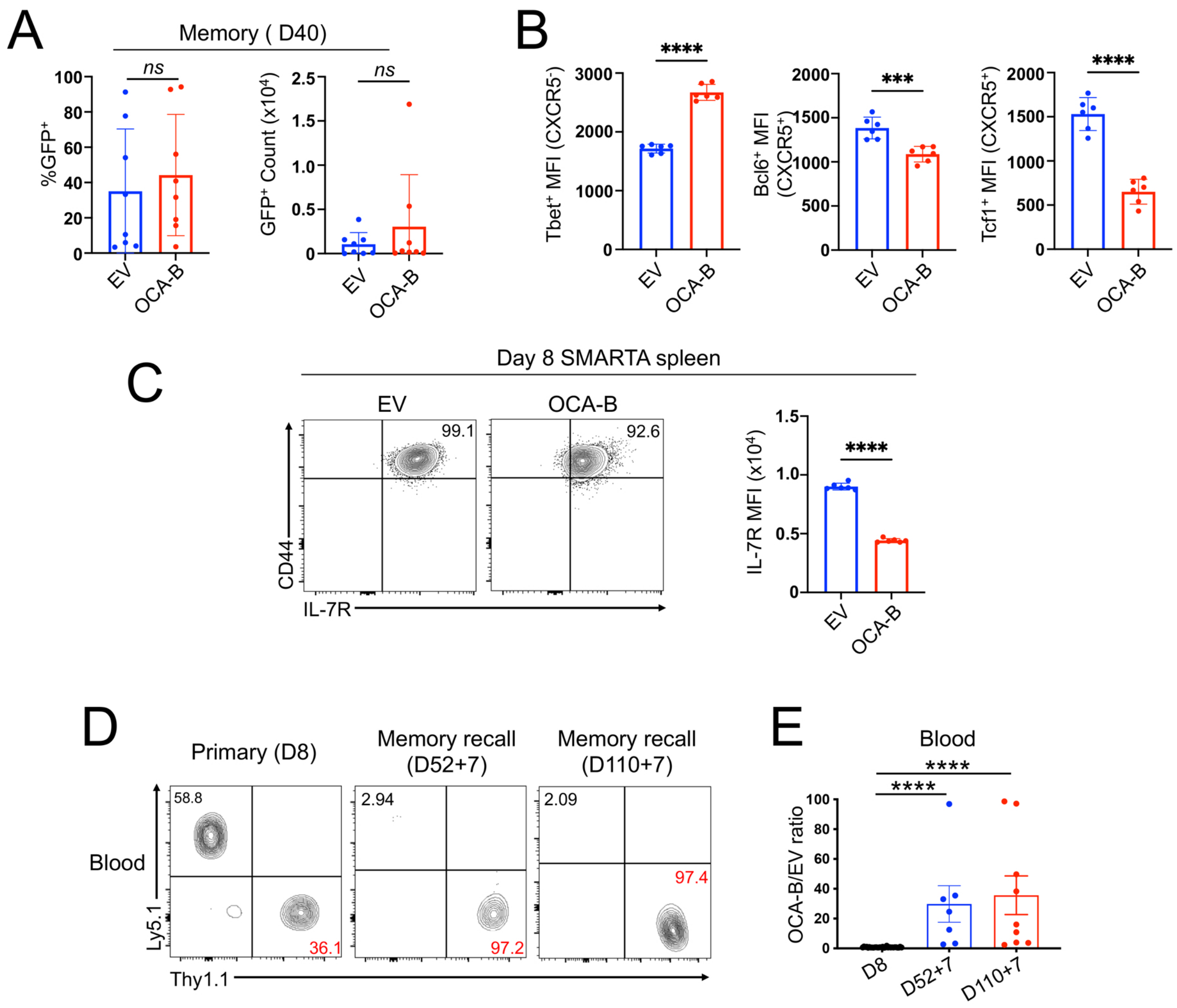
A
Memory ( D40)
ns
%GFP+
EV
OCA-B
GFP+ Count (x10^4)
B
****
Tbet+ MFI (CXCR5-)
***
Bcl6+ MFI (CXCR5+)
Tcf1+ MFI (CXCR5+)
C
Day 8 SMARTA spleen
99.1
92.6
CD44
IL-7R
IL-7R MFI (x10^4)
D
Primary (D8)
Memory recall (D52+7)
Memory recall (D110+7)
Blood
58.8
36.1
2.94
97.2
2.09
97.4
Ly5.1
Thy1.1
E
OCA-B/EV ratio
D8
D52+7
D110+7

**Fig. S2.** Additional markers and quantifications collected from SMARTA cells transduced with empty vector or OCA-B. (*A*) Related to Fig. 1*D* except splenic cells from static memory timepoint animals (D40) were quantified by %GFP and GFP counts. n=8 for all quantifications. (*B*) Related to Fig. 1*F*, *G* except additional quantification is shown for mean fluorescence intensities: Tbet MFI of $CXCR5^-$ cells (left chart), Bcl6 MFI of $CXCR5^+$ cells (center chart) and TCF1 MFI of $CXCR5^+$ cells (right chart). n=6 for all quantifications. (*C*) Related to Fig. 1*F*, *G* except additional staining and quantification is shown for IL-7R. n=6. (*D*) Related to Fig. 1*J* except additional staining is shown for peripheral blood. (*E*) Related to Fig 1*K* except additional quantification is shown for peripheral blood. All mice were bled at D8. Remaining mice were bled at subsequent timepoints.

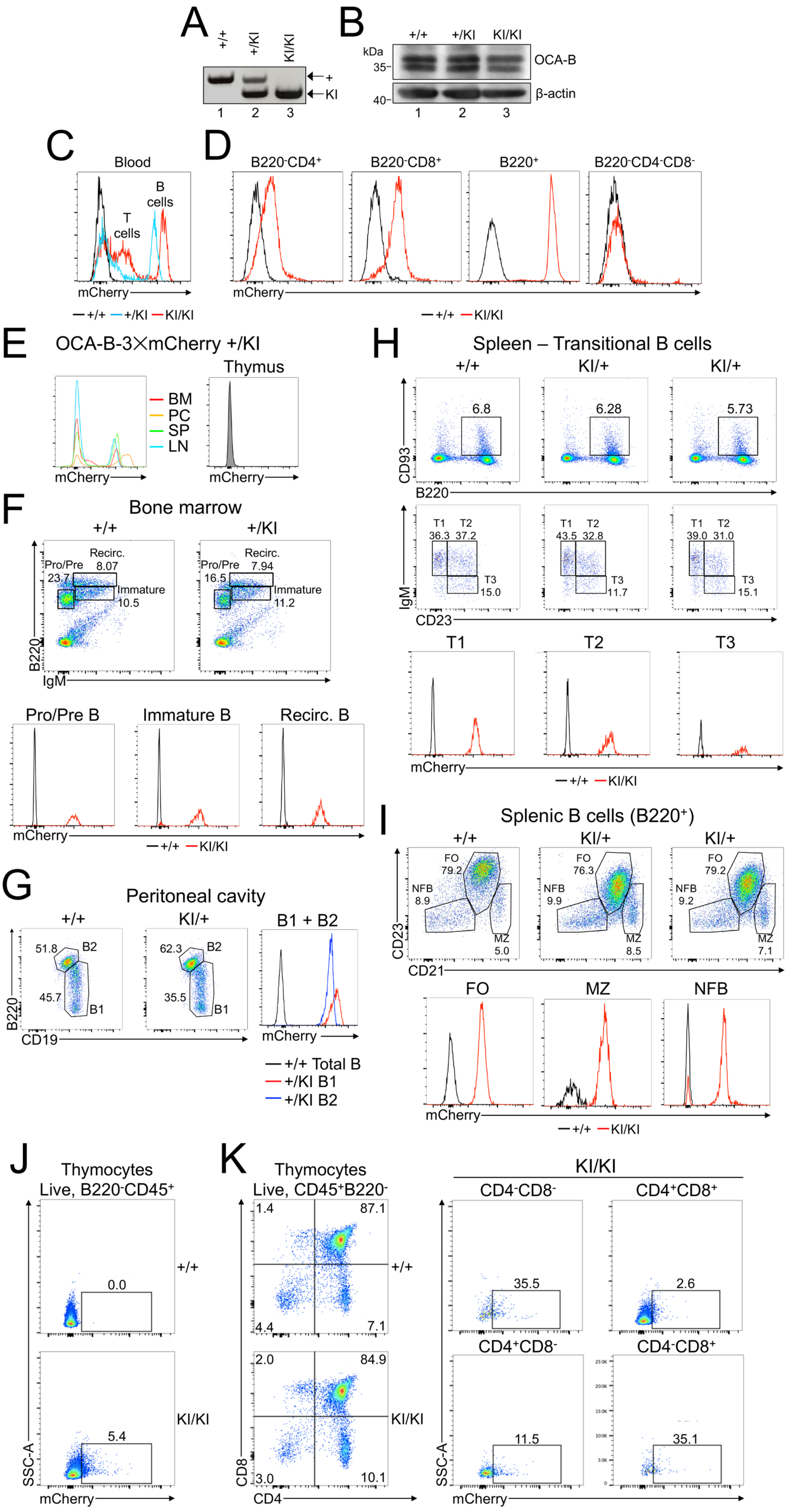
A
+/+
+/KI
KI/KI
+
KI
1
2
3
B
+/+
+/KI
KI/KI
kDa
35
40
OCA-B
β-actin
C
Blood
T cells
B cells
mCherry
+/+
+/KI
KI/KI
D
B220-CD4+
B220-CD8+
B220+
B220-CD4-CD8-
E
OCA-B-3×mCherry +/KI
Thymus
BM
PC
SP
LN
F
Bone marrow
Pro/Pre B
Immature B
Recirc. B
G
Peritoneal cavity
B1 + B2
+/+ Total B
+/KI B1
+/KI B2
H
Spleen – Transitional B cells
T1
T2
T3
I
Splenic B cells (B220+)
FO
MZ
NFB
J
Thymocytes
Live, B220-CD45+
K
Thymocytes
Live, CD45+B220-
KI/KI
CD4-CD8-
CD4+CD8+
CD4+CD8-
CD4-CD8+
SSC-A
mCherry
CD8
CD4

**Fig. S3.** OCA-B-3×mCherry knock-in reporter mouse characterization in B and developing T cell populations. (*A*) PCR genotyping OCA-B-3×mCherry reporter mice. PCR yields a 494 bp product for the wild-type *Pou2af1* allele and a 373 bp product for the targeted *Pou2af1* allele. +=wild-type allele. KI=knock-in allele. (*B*) OCA-B immunoblot using splenocytes from wild-type (+/+) and heterozygous (+/KI) or homozygous knock-in OCA-B-3×mCherry knock-in mice. β-actin was used as a loading control. (*C*) Flow cytometry showing mCherry expression in peripheral blood cells of wild-type mice (+/+), or either heterozygous (+/KI) or homozygous (KI/KI) OCAB-3×mCherry knock-in reporter mice. (*D*) Similar flow cytometry plots as in (C) except gated on the indicated populations. (*E*) mCherry (OCA-B) expression in a 4.5 week-old heterozygous female OCA-B-3×mCherry reporter mouse. Cells were harvested from bone marrow (BM), peritoneal cavity (PC), spleen (SP), lymph node (LN) or thymus. After removal of red blood cells, mCherry expression was measured in total cell populations. (*F*) Analysis of bone marrow B cell populations from +/+ and +/KI reporter mice. Cells were stratified into pro/pre, immature and recirculating B cell compartments based on B220 and IgM expression. Top plots show relative proportions of the cells in mice of the indicated genotypes. Bottom plots show mCherry levels in the gated populations. (*G*) Wild-type (+/+) and heterozygous knock-in (+/KI) peritoneal cavity B cells were further stratified into B1 and B2 populations based on B220 and CD19 expression. Top plots show relative proportions of B1 ($B220^{lo}CD19^{hi}$) and B2 ($B220^{hi}CD19^{lo}$) cells in wild-type and two example heterozygous reporter mice. Right plots show mCherry expression in gated populations. (*H*) Splenic transitional B cells were stratified using CD93 (AA4.1), CD23 and IgM antibodies. One wild-type and two example heterozygous reporter mice are shown. (*I*) Similar analysis to (H) except using CD21 and CD23 antibodies to visualize follicular and MZ B cell populations. (*J*) mCherry expression in total B220-negative thymocytes from wild-type (+/+) or homozygous knock-in reporter mice was analyzed by flow cytometry. Specific pathogen-free mice were used. (*K*) Thymocytes were stained with CD4 and CD8 antibodies to visualize double-negative, double-positive and single-positive T cell developmental stages in homozygous knock-in reporter mice. Right: mCherry expression in developing reporter T cells was subdivided by quadrant.

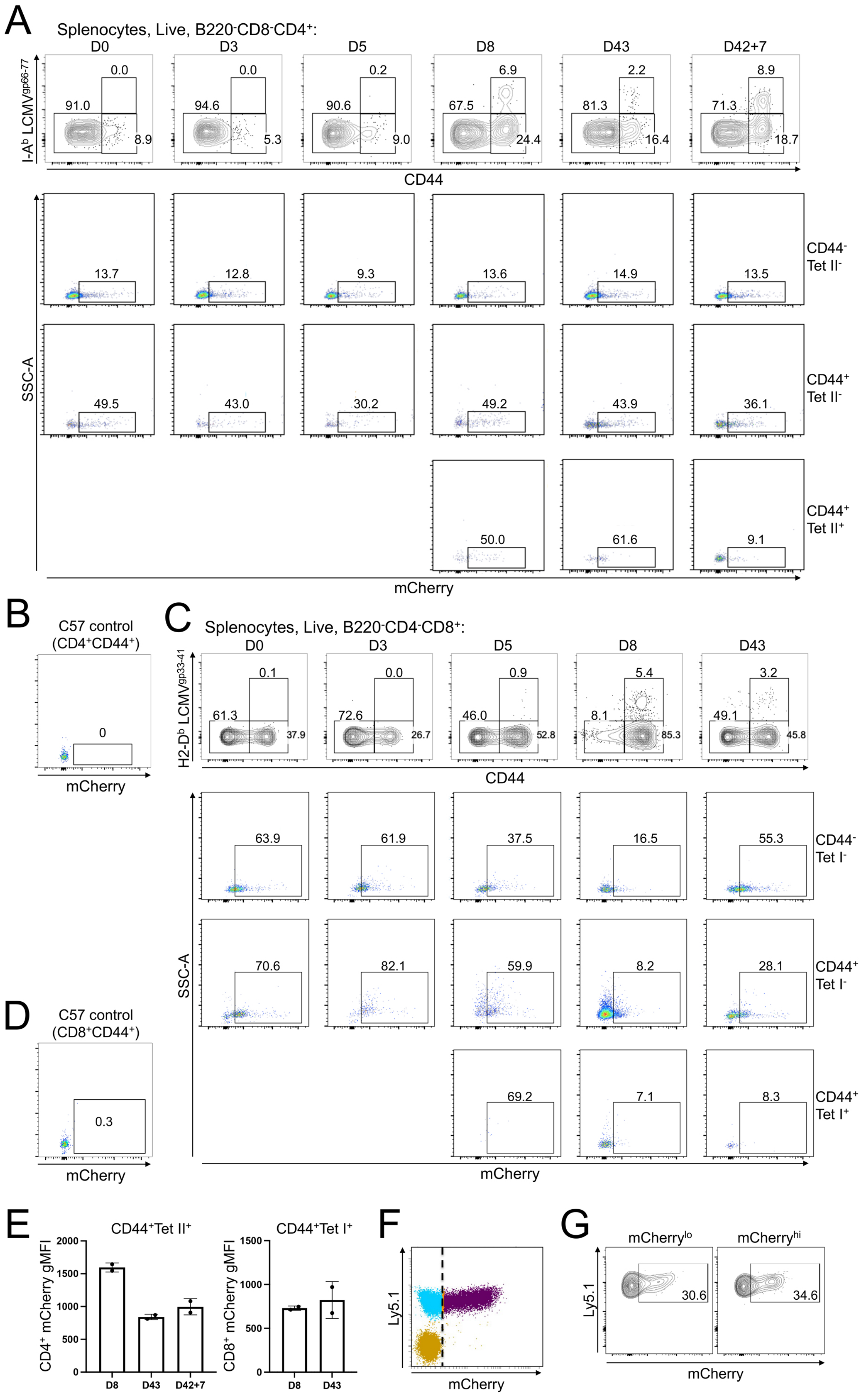
A
Splenocytes, Live, B220-CD8-CD4+:
D0
D3
D5
D8
D43
D42+7
I-Ab LCMVgp66-77
0.0
91.0
8.9
0.0
94.6
5.3
0.2
90.6
9.0
6.9
67.5
24.4
2.2
81.3
16.4
8.9
71.3
18.7
CD44
13.7
12.8
9.3
13.6
14.9
13.5
CD44- Tet II-
SSC-A
49.5
43.0
30.2
49.2
43.9
36.1
CD44+ Tet II-
50.0
61.6
9.1
CD44+ Tet II+
mCherry
B
C57 control (CD4+CD44+)
0
mCherry
C
Splenocytes, Live, B220-CD4-CD8+:
D0
D3
D5
D8
D43
H2-Db LCMVgp33-41
0.1
61.3
37.9
0.0
72.6
26.7
0.9
46.0
52.8
5.4
8.1
85.3
3.2
49.1
45.8
CD44
63.9
61.9
37.5
16.5
55.3
CD44- Tet I-
SSC-A
70.6
82.1
59.9
8.2
28.1
CD44+ Tet I-
69.2
7.1
8.3
CD44+ Tet I+
mCherry
D
C57 control (CD8+CD44+)
0.3
mCherry
E
CD44+Tet II+
CD4+ mCherry gMFI
2000
1500
1000
500
0
D8
D43
D42+7
CD44+Tet I+
CD8+ mCherry gMFI
1500
1000
500
0
D8
D43
F
Ly5.1
mCherry
G
mCherrylo
mCherryhi
Ly5.1
30.6
34.6
mCherry

**Fig. S4.** OCA-B expression in antigen-responsive T cells following LCMV infection. (*A*) Flow cytometry plots are shown of homozygous OCA-B-3×mCherry knock-in reporter mice at different timepoints following LCMV infection (D0-43) and following rechallenge with Lm-gp61 (D42+7). Mice were challenged on different days and spleens were harvested and processed on the same day. Top panels: percentages of cells expressing CD44 and class II LCMV tetramer (I-$A^b$ $LCMV^{gp66-77}$) are shown. Bottom panels: for the same timepoints cells gated from each quadrant in the top panels are plotted. Each flow panel is representative of biological duplicate mice sacrificed at each timepoint. (*B*) Parallel gating for a control C57BL/6 mouse lacking the reporter allele. (*C*) Cells from the same mice as above were also stained with class I LCMV tetramers (H2-$D^b$ $LCMV^{gp33-41}$). Analysis of recall response timepoints was not possible for antigen-specific CD8 cells because mice were rechallenged with Lm-gp61, which lacks CD8 immunodominant epitopes. (*D*) Parallel gating for a C57BL/6 mouse lacking the reporter allele. (*E*) Geometric mean fluorescence intensity (gMFI) of $CD4^+$ and $CD8^+$ cells falling within the $mCherry^+$ gate was quantified for n=2 mice and averaged. (*F*) Related to Fig. 5*A-D* showing the mCherry gating strategy used to identify $mCherry^{hi}$ and $mCherry^{lo}$ cells. Nonfluorescent, Ly5.$1^{neg}$ host cells (brown) provided an internal standard for lack of mCherry expression. (*G*) Related to Fig. 5*H* except showing a representative flow plot of adoptively transferred $mCherry^{hi}$ and $mCherry^{lo}$ cells after rechallenge.
